# *Drosophila* have distinct activity-gated pathways that mediate attraction and aversion to CO_2_

**DOI:** 10.1101/227991

**Authors:** Floris van Breugel, Ainul Huda, Michael H. Dickinson

## Abstract

Carbon dioxide is a volatile and broad signal of many organic processes, and serves as a convenient cue for insects in search of blood hosts^1–6^, flowers^7^, decaying matter^8–11^, communal nests^12^, fruit^13^, and wildfires^14^. Curiously, although *Drosophila melanogaster* feed on yeast that produce CO_2_ and ethanol during fermentation, laboratory experiments suggest that flies actively avoid CO_2_^15–25^. Here, we resolve this paradox by showing that both flying and walking fruit flies do actually find CO2 attractive, but only when they are in an active state associated with foraging. Aversion at low activity levels may be an adaptation to avoid CO_2_-seeking-parasites, or succumbing to respiratory acidosis in the presence of high concentrations of CO_2_ that are occasionally found in nature^26,27^. In contrast to CO_2_, flies are attracted to ethanol in all behavioral states, and invest twice as much time searching near ethanol compared to CO_2_. These behavioral differences reflect the fact that whereas CO_2_ is a generated by many natural processes, ethanol is a unique signature of yeast fermentation. Using genetic tools, we determined that the evolutionarily ancient ionotropic co-receptor IR25a is required for both CO_2_ and ethanol attraction, and that the receptors previously identified for CO_2_ avoidance are not involved. Our study lays the foundation for future research to determine the neural circuits underlying both state- and odorant-dependent decision making in *Drosophila*.

---

The life history of the fruit fly, *Drosophila melanogaster*, revolves around fermenting fruits, where they feed, mate, and deposit eggs. Their lifecycle from egg to adult takes approximately 10-14 days, roughly the same amount of time that most ripe fruit takes to decay. Thus, upon emerging from their puparia, adult flies need to locate a fresh ferment. The primary compounds produced by yeast fermentation are ethanol and CO_2_. Because of its high volatility, CO_2_ emission is greatest near the start of fermentation, whereas the ethanol emission increases more slowly (Fig. 1a). Other odors associated with fermentation, such as acetic acid and ethyl acetate, form later when bacteria begin to break down the ethanol.

**Figure 1.**
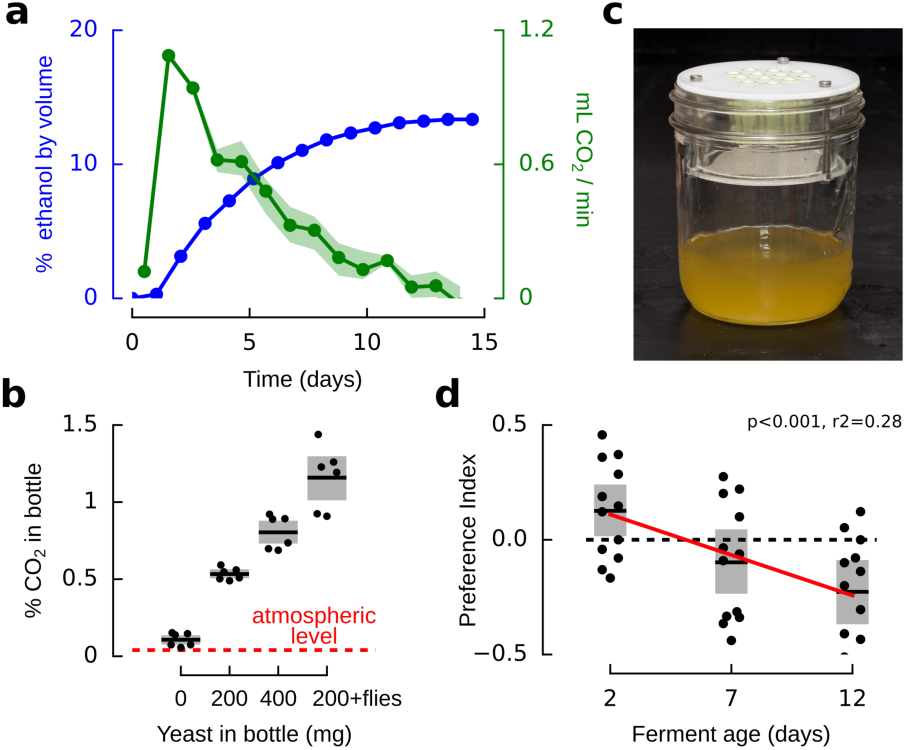
*Drosophila* prefer early fermentations, at peak CO_2_ production. **a,** Alcohol by volume for apple juice and sugar fermented with champagne yeast over the course of 2 weeks, measured with a hydrometer. CO_2_ production was calculated from the stoichiometry of fermentation (1 sugar molecule yields 2 ethanol + 2 CO_2_), corresponding to the derivative of alcohol by volume. **b,** CO_2_ concentration in 500 mL fly rearing bottles under common laboratory conditions. See Fig. S1 for methods and calibration details. **c,** Trap assay. **d,** Preference index exhibited by flies in three 2-choice assays, using traps shown in **b.** Flies were presented with two traps: one was a completed 14 day old ferment which had been stored in the refrigerator, the second was a fresh ferment aged 2, 7, or 12 days old. Positive preference index indicates a preference for the fresh ferment. Mean and standard deviation of total captured flies for each trial: 105±59. In all panels, shading indicates bootstrapped 95% confidence intervals.

To find an active rot, flies should therefore search near sources of both CO_2_ and ethanol. Variable air currents make it difficult to estimate the exact concentration of CO_2_ emitted from ferments in the wild; however, we measured the CO_2_ concentration in 500 mL bottles used to rear flies in many laboratories (see Methods and Fig. S1). Such bottles contain 0.5-1% CO_2_ depending on the amount of yeast and flies present (Fig. 1b), and serve as effective traps if left without a lid. In trap assays (Fig. 1c), *Drosophila* showed a preference for 2-day-old apple juice ferments compared to older solutions in which the yeast had flocculated and were no longer producing CO_2_ (Fig. 1d).

This casual evidence that CO_2_ attracts *Drosophila* contradicts many prior studies that concluded flies actively avoid CO_2_ in small chambers and T-mazes^15–25^. To study how flies respond to different odors under more ethologically relevant conditions, we recorded the flight trajectories^28,29^ of flies in a wind tunnel containing a fruit-sized landing platform, which we programmed to periodically release plumes of CO_2_ or ethanol (Fig. 2a-b). In the presence of either odor, flies were far more likely to approach and land on the platform. They also approached a dark spot on the floor of the wind tunnel (Fig. 2c-d), consistent with prior experiments with flies and mosquitoes^2,29^. Flies were more likely to approach the platform or the dark spot in the presence of ethanol compared to CO_2_, but were equally likely to land in the presence of either odor (Fig. 2e).

**Figure 2.**
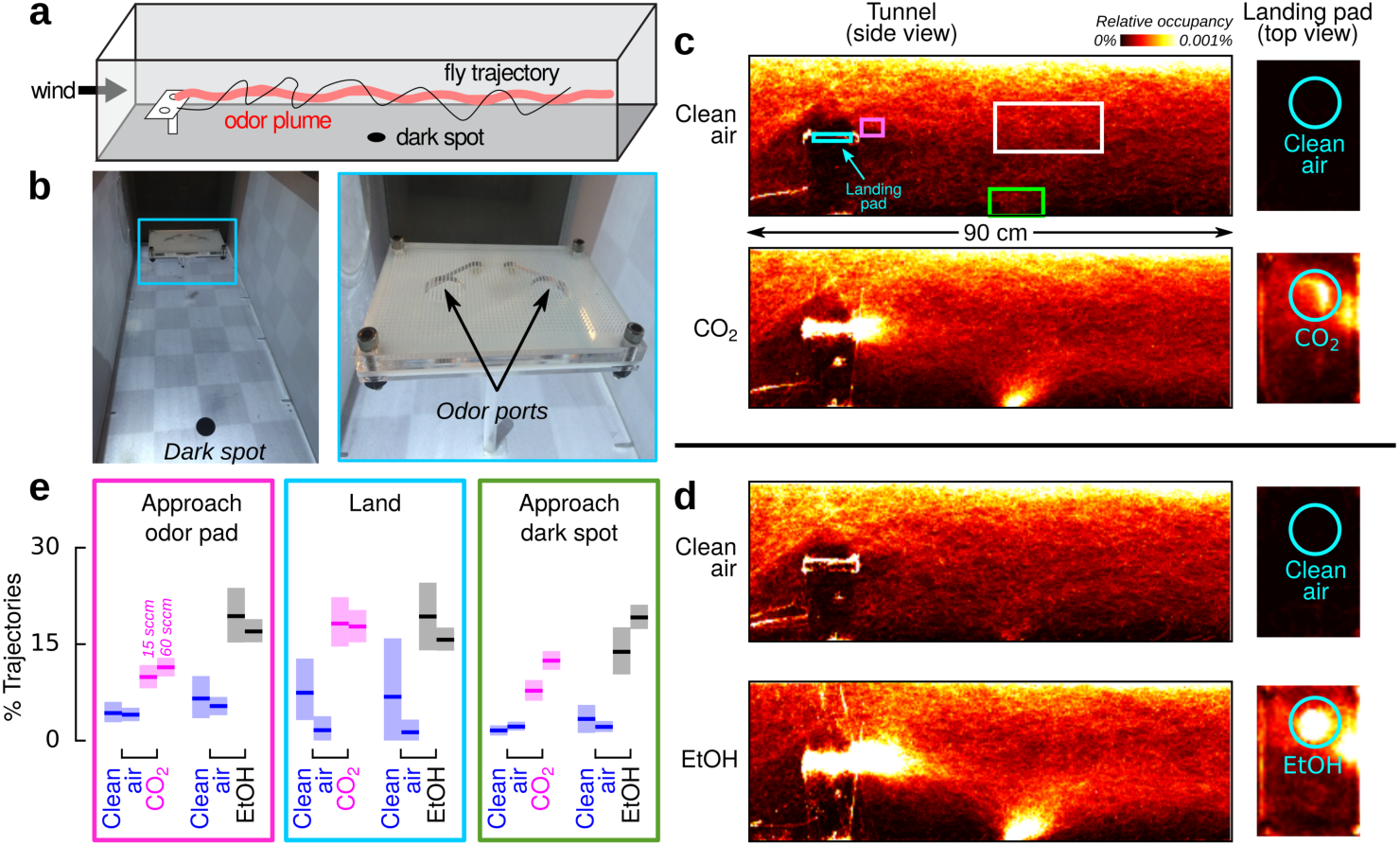
*Drosophila* are attracted to both ethanol and CO_2_ in flight. **a,** Diagram of our wind tunnel, illustrating the relative position of the odor platform and a conspicuous visual feature. **b,** Photograph of the wind tunnel, viewing upwind, and close up of the odor-emitting landing platform. **c, d** Heat-maps indicating relative occupancy of flies in the presence of either CO_2_ or ethanol. Cohorts of 12 flies were introduced into the wind tunnel and their behavior recorded over the course of 16 hrs. Throughout the experiment, 100 sccm of clean air emerged from both odor ports. For 30 min every hour, 60 sccm of either CO_2_ or clean air bubbled through pure ethanol was added to one of the odor ports. Control data come from the 30 min segments of clean air prior to each odor stimulus. Number of cohorts: 9 (CO_2_), 6 (ethanol). Number of trajectories total: 59,900 – 101,000 per panel. e, Fraction of trajectories from c and d that enter one of the colored volumes in c relative to another volume. Approaches to landing pad: magenta/white; landings: cyan/magenta; approaches to dark spot: green/white. Number of trajectories per condition: 44-1288 (control), 228-1815 (odor). Experiments were performed on two different flowrates (i.e. concentrations): 15 sccm and 60 sccm. In all panels, shading indicates bootstrapped 95% confidence intervals.

To quantify the behavior of flies after they land, we designed a new platform suitable for automated tracking (Fig. 3a-b). For a flow rate of 60 sccm CO_2_, the CO_2_ concentration near the surface of the platform was approximately 3% (Fig. 3c, S2). After landing near a source of CO_2_, ethanol, or apple cider vinegar, flies exhibited local search behavior (Fig. 3d), which we summarized using four descriptive statistics (Fig. 3e, S3). Flies spent approximately twice as much time exploring the platform in the presence of ethanol compared to CO_2_ or any other odor. Vinegar elicited smaller local searches than either CO_2_ or ethanol. While searching on the platform, flies approached the odor source most frequently for ethanol and CO_2_. Vinegar elicited slightly fewer approaches compared to CO_2_, consistent with the hypothesis that vinegar might indicate a less favorable, late-stage ferment. Flies spent significantly less time standing still on the platform in the presence of CO_2_ compared to any other odor, exhibiting an overall mean walking speed greater than 2mm s^−1^. When combined in a single odor stream, CO_2_ and ethanol together elicited a stronger search behavior than that exhibited to either odor alone.

**Figure 3.**
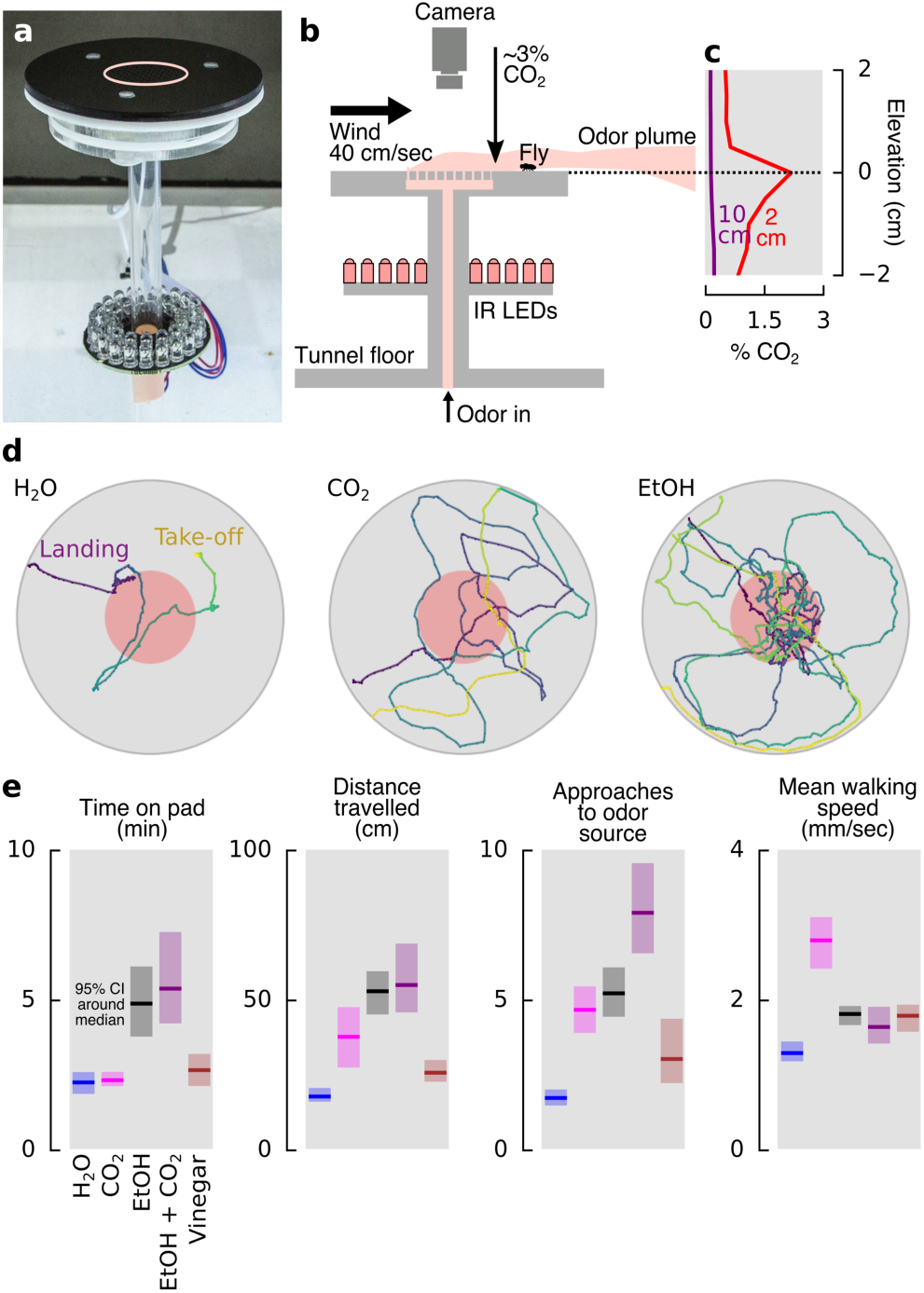
*Drosophila* are attracted to CO_2_ while walking, but spends twice as much time exploring ethanol sources. **a,** Photograph of landing platform for observing walking behavior. **b,** Cross-sectional diagram of the odor platform. **c,** CO_2_ concentration profile for two altitude transects 2 cm and 10 cm downwind from the platform at a 60 sccm flow rate (see supplemental materials, Fig. S2). **d,** Stereotypical trajectories in response to odors. Color encodes time. **e,** Four descriptive statistics summarizing flies’ behavior on the platform in response to different odors. We used a flow rate of 60 sccm for each odor, except for the ethanol + CO_2_ combination which consisted of 60 sccm clean air bubbled through ethanol with 15 sccm of CO_2_ added. See Fig. S3 for experiments done at additional flow rates. Horizontal bars: median value; shading: 95% confidence interval. Number of trajectories for each case is between 121 and 193.

One prior study using a tethered flight assay showed that *Drosophila* are attracted to CO_2_ while flying, a result that was attributed to the influence of the elevated levels of octopamine during flight^30^. Our results confirm this observation in freely-flying flies; however, we also found that flies continue to be attracted to CO_2_ after they land. One possible explanation for this discrepancy is that the elevated levels of octopamine during flight might influence the flies’ reactions to CO_2_ for a short time after landing. To test this hypothesis, we built an enclosed walking arena in which flies were unable to fly (Fig. 4a, S4-6), and presented them with pulses of 5% CO_2_ (close to the 3% concentration that elicited attraction in the wind tunnel assay). Starved flies presented with CO_2_ after acclimating to the arena for 10 min exhibited aversion, as has been previously reported in such chambers (Fig. 4b). However, if allowed to acclimate for two hours and then given a pulse of CO_2_, the animals exhibited attraction (Fig. 4c).

**Figure 4.**
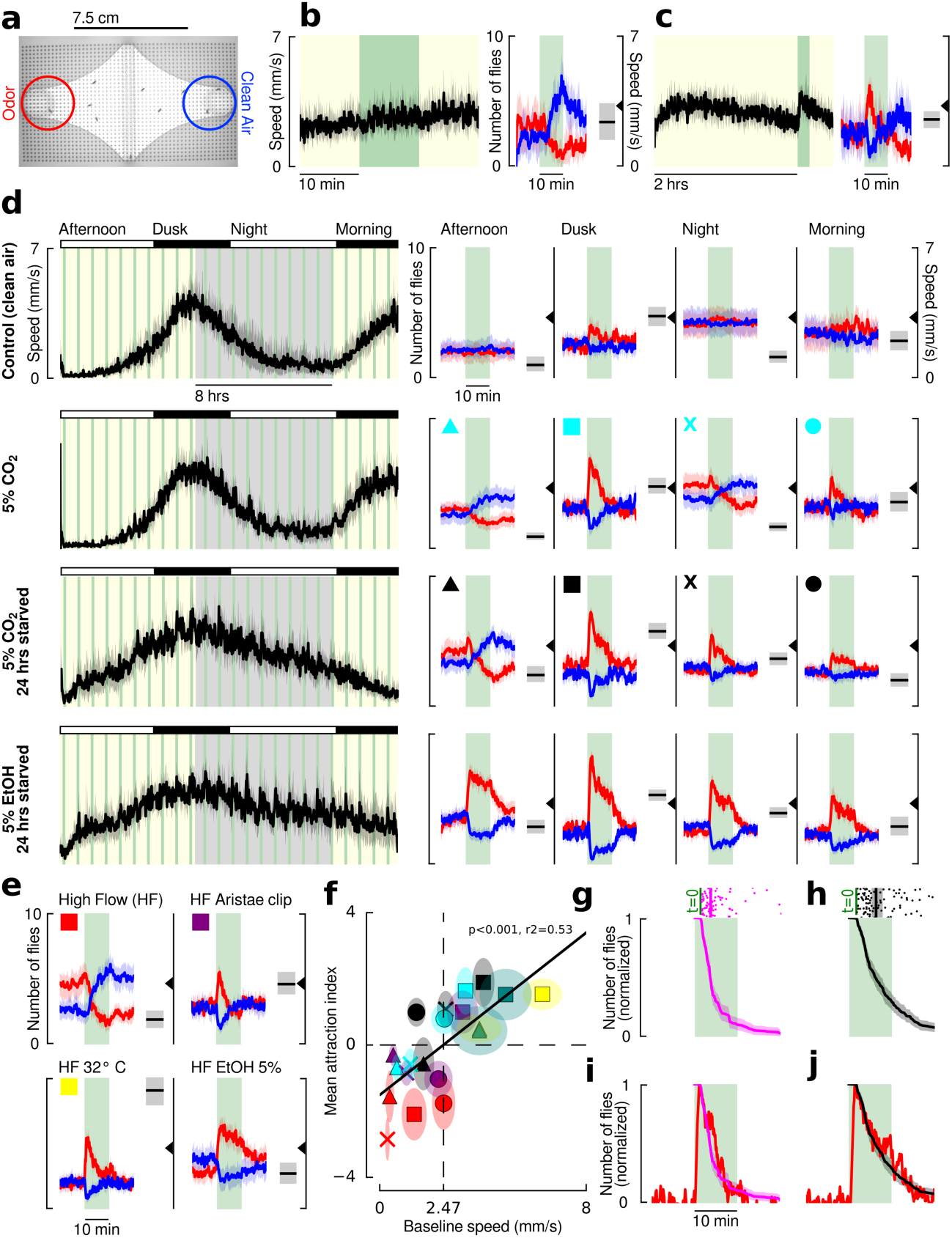
*Drosophila* are briefly attracted to CO_2_ during periods of high activity, in contrast to their activity independent and sustained attraction to ethanol. **a,** Image of walking arena, with blue and red regions of interest (ROI) for counting flies near clean air and odor. **b,** (left) Mean speed of 10 starved flies over 30 min. Green indicates times when 1 sccm of odor (or clean air control) was added to the constant humidified clean air bulk flow (20 sccm) from alternating sides. (right) Number of flies near the CO_2_ (red) and clean air (blue) over time. Black bar and shading shows the mean and 95% confidence interval of the flies’ speeds five min prior to the odor presentation. The triangle provides a reference point. N=8 cohorts of 10 flies. **c,** Same as **b,** but with a 2-hour acclimatization period prior to the CO_2_ presentation. N=10 cohorts of 10 flies each. Experiments for b and c were both done during flies’ peak activity time (dusk). d, (left) Mean speed of flies for a 20 hr experiment. Yellow/gray indicate entrained day/night cycle (experiments are done in darkness). (right) Data plotted as in b, for four different time frames. **e,** Data for the dusk time frame of experiments done under 100 sccm bulk flow conditions. In these experiments 5 sccm odor was added to achieve the same concentration as in e. Experiments were performed with 5% CO_2_ stimuli on in-tact flies (red), flies with aristae surgically removed (purple), and in-tact flies at 32° under a heat lamp (yellow). Finally, we performed experiments with in-tact flies in response to ethanol. **f,** Summary of CO_2_ responses presented in d and e (color and shape encodes experiment and time of day). Green data points are for experiments done at 20 sccm bulk flow at 32° C. Mean attraction index is calculated as the mean number of flies near the CO_2_ over the 10-min presentation, minus the number of flies near the CO_2_ 5 min prior to the presentation. Baseline speed refers to the mean speed of all the flies 5 min prior to the CO_2_ presentation. g, Scattergram shows the amount of time each fly spent searching the odor platform in the wind tunnel from Fig. 3a in the presence of 60 sccm CO_2_ (data is repeated from Fig. 3e). Time trace is the bootstrapped mean and 95% confidence intervals for the normalized number of flies that would have been on the platform had all the flies landed simultaneously. The green shading is only provided for reference – the odor never turned off in these wind tunnel experiments. **h,** Same as **g,** but for ethanol. **i,** Time trace from **g** overlaid on the normalized number of un-starved flies near the 5% CO_2_ source during the dusk time period in the walking arena, copied from panel d. **j**, Time trace from **h** overlaid on the normalized number of un-starved flies near the 5% ethanol source during the dusk time period in the walking arena, data not shown in **d.** We chose un-starved flies for the comparisons because wind tunnel experiments were done with un-starved flies. We chose the 60 sccm case for the comparison because the CO_2_ concentration in the wind tunnel matches the 5% CO_2_ stimulus given in the walking experiments. Throughout the figure, shading around data indicates 95% confidence intervals. All experimental combinations were performed with 6 cohorts of 10 flies each.

To study their responses to CO_2_ in more detail, we recorded the behavior of flies for 20 continuous hours in darkness, while offering 10 min long presentations of CO_2_ from alternating sides of the arena every 40 minutes (Fig. 4d). Throughout the experiments, both sides of the arena received 20 sccm of air saturated with water vapor. The flies exhibited a clear circadian rhythm in their activity within the chamber, as indicated by their mean walking speed. At times of peak activity — near their entrained dusk and dawn — flies showed a strong initial attraction to CO_2_, which decayed stereotypically during the 10 min presentation. At times of low activity — at mid-day and during the night — the flies exhibited a mild aversion to CO_2_. Starving flies for 24 hours prior to placing them in the chamber (instead of just 3 hours) changed their activity profile, resulting in a slightly elevated attraction during their subjective night. Ethanol, in contrast, elicited sustained attraction regardless of baseline activity or time of day (Fig. 4d).

Our experiments thus far suggest a possible correlation between activity and attraction to CO_2_. To test this hypothesis, we made several other environmental manipulations that are known to alter activity: increased temperature and wind speed (Fig. 4e). When we increased our bulk flow rate to 100 sccm, flies exhibited a peak walking speed (at dusk) of about 1.5 mm s^−1^, nearly half the speed we measured when using a flow rate of 20 sccm. This result is consistent with observations that flies stop moving in the presence of wind^31^. Instead of showing attraction, these flies exhibited aversion to 5% CO_2_ when it was presented at this higher flow rate; however, they still exhibited attraction to ethanol (Fig. 4e). This result helps to explain why previous studies that used high bulk airflow rates of 100-1000 sccm to present CO_2_^16,24^ observed aversion. To further explore the effect of wind speed on behavior, we clipped the flies’ aristae. This manipulation destroys their primary means of detecting airflow but does not interfere with the detection of odors^32^. The aristae-less flies exhibited the same walking speed and attraction to CO_2_ at the high flow rate as exhibited by normal flies at the low flow rate. We also warmed flies with intact aristae to 32° C, which increased their baseline activity. These flies also exhibited attraction to CO_2_ at the higher flow rate. Pooling data across all our experimental conditions, we found that flies were attracted to CO_2_ when they had a baseline walking speed above ~2.4 mm s^−1^ (Fig. 4f). This result is similar to the mean walking speed value we observed in our wind tunnel assay, which was higher for CO_2_ than the other odors we tested. This suggests that there may be some underlying physiological connection between circuits regulating locomotor activity and those regulating CO_2_ attraction. To confirm that activity dependent attraction to CO_2_ is not a function of social interactions, we also performed experiments on 30 single flies, which on average behaved exactly as the cohorts of 10 (Fig. S7). We also tested three concentrations of CO_2_ (1.7%, 5%, 15%) and found that 5% elicited the strongest response, consistent with our wind tunnel experiments (Fig. S8).

Although flies’ responses to ethanol and CO_2_ were similar during the first minute of the stimulus, the attraction to ethanol was more sustained. The time course of behavior was remarkably similar in the walking arena and wind tunnel (Fig. 4g-j), suggesting that the behavioral dynamics of olfactory attraction are robust to the stimulus environment, and may represent an adaptation for utilizing information that ecologically broad (CO_2_) and more specific (ethanol) odorants provide.

Despite the ethological importance of both ethanol and CO_2_ as food cues for *Drosophila*, the olfactory receptors used to detect these odors during foraging are not known. To determine if CO_2_ attraction is mediated by either an olfactory (OR) or ionotropic (IR) receptor, we used our apparatus to test an IR8a; IR25a; Orco, Gr63a quadruple mutant, which lack the OR and IR co-receptors as well as a CO_2_-sensitive gustatory receptor (Fig. 5a-b). These near-anosmic mutants exhibited no detectable behavioral response to CO_2_. Flies in which we surgically removed the 3^rd^ antennal segment also showed no response to CO_2_, despite otherwise normal levels of activity. Together with our arista ablations (Fig. 4e), these experiments show that CO_2_ attraction is mediated by the olfactory system in the 3^rd^ antennal segment.

**Figure 5.**
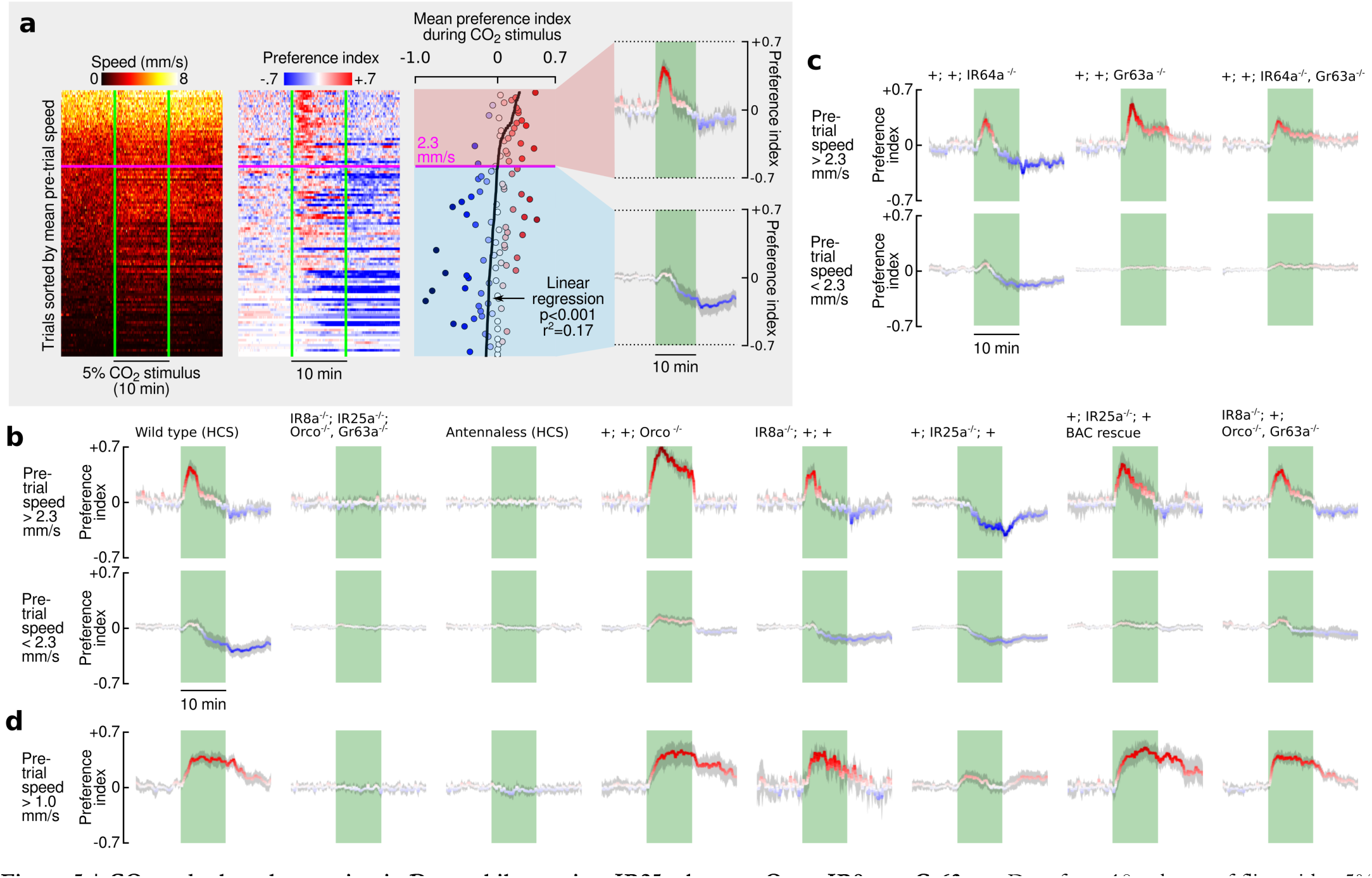
CO_2_ and ethanol attraction in Drosophila requires IR25a, but not Orco, IR8a, or Gr63a. **a,** Data from 10 cohorts of flies with a 5% CO_2_ stimulus sorted by the mean speed (S) during the reference period of 5 min prior to the odor stimulus (*S̅*|_*Ref*_). To achieve a wide range of baseline activities, 4 of the cohorts were starved for 24 hrs, 3 were starved for 3 hrs, and 3 were starved for 3 hrs and heated to 32° C. Preference index (PI) was calculated in two steps: (1) PI_0_ = (N_odor_ – N_control_)/N_total_; (2) *PI* = *PI*_0_ – *PI*_0_̅|_*Ref*_. Where N denotes thhe number of flies, and N_total_=10. Next, we calculated the mean PI during the stimulus period and determined the linear regression with respect to *S̅*|_*Ref*_, and used the intercept to cluster the data into two groups: high activity and low activity. For these two groups, we calculated the mean PI over time. **b-c,** Data plotted as in last panel of **a,** for different manipulations and mutants, using the intercept found in a of 2.3 mm/sec to cluster the data. All flies were presented with randomly interleaved stimuli of 0% or 5%; only responses to 5% are shown here. Responses to 0% stimuli are shown in Fig. S9. IR25a BAC flies did not respond to CO_2_ when heated; only the unheated data is shown here (see Fig. S10). **d,** Same as **b,** but for ethanol odors. Since ethanol attraction is not activity dependent, we used 5 cohorts of 24 hr starved flies, and used a mean speed of 1 mm/sec to cluster the data. Only the higher activity is shown; the low activity flies show small responses ethanol. Shading indicates bootstrapped 95% confidence intervals for all panels.

Prior research has shown that flies’ aversion to CO_2_ is mediated by a pair of olfactory receptors, Gr63a and Gr21a^15,18,33^, with high concentrations of CO_2_ also being detected by the acid-sensitive ionotropic receptor IR64a^20^ (which operates together with the co-receptor IR8a). Mutant flies lacking the IR64a receptor showed no significant change in their behavior compared to wild type (Fig. 5c). Mutants lacking the Gr63a receptor exhibited no aversion to CO_2_ (Fig. 5c), consistent with the prior literature; however, the same animals were still attracted to CO_2_ when more active. Homozygous Gr63a/IR64a double mutants behaved similarly to the Gr63a mutants. It is noteworthy that the characteristic decaying time course of attraction was unaffected in Gr63a mutants, even though these flies showed no aversion. This suggests that the decay in attraction to CO_2_ is not caused by an increase in aversion over time.

Collectively, our results suggest that none of the canonical CO_2_ receptors are responsible for attraction. The OR class of receptors is unlikely to mediate the response; indeed, Orco mutants exhibited a sustained attraction to CO_2_. IR8a mutants also exhibit normal attraction to CO_2_. Ir25a mutants, however, exhibited only aversion to CO_2_ at all activity levels, whereas rescuing Ir25a with a bacterial artificial chromosome^34^ rescued their attraction. Mutant flies lacking Orco, Ir8a, and Gr63a exhibit wild type attraction to CO_2_, indicating that none of these proteins are necessary for CO_2_ attraction. We further verified our results by testing a different mutant allele of the IR25a, which behaved in the same manner (Fig. S10).

Although ethanol is the most ethologically relevant cue related to fermentation and elicits the strongest attraction, the receptors for ethanol are still not known. We used the same set of co-receptor mutants to determine that IR25a, but not Orco or IR8a, is required for attraction to ethanol (Fig. 5d). IR25a is, however, not required for attraction to other odors associated with foraging, like apple cider vinegar (Fig. S9), confirming that these mutants are still capable of exhibiting attractive behaviors.

Prior studies reporting aversion to CO_2_ have suggested that it serves as a pheromonal cue *(Drosophila* Stress Odor, DSO) by which stressed flies signal others to flee a local enviroment^15^. Our result that active flies are attracted to CO_2_ is not consistent with this hypothesis. An alternative explanation for the prior findings is that stressed insects release CO_2_ simply because they have it stored in their tracheal system as part of the normal process of discontinuous respiration^35,36^. Indeed, we found that even mosquitoes (which are strongly attracted to CO_2_) release CO_2_ when shaken (Fig. S12). We suggest that the DSO hypothesis is a by-product of two unrelated behaviors: the release of tracheal CO_2_ by agitated flies and the avoidance of CO_2_ while in a behavioral state related to either low activity levels or being recently introduced to a new chamber (and thus likely to be in a behavioral state more associated with exploring a new environment than foraging). This aversive behavior may be an adaptation that helps sleeping flies either minimize encounters with parasites that are themselves attracted to CO_2_ as a means of finding hosts (parasitic wasps of *Drosophila* are attracted to yeast products^37^ and thus likely CO_2_; other hematophagous parasites are often attracted to CO_2_^1,3,4,6^), or avoid succumbing to respiratory acidosis in the presence of high concentrations of CO_2_. Examples of insects being fatally attracted to high levels of CO_2_ have been reported in the literature^27^, and we have replicated this behavior in the lab (Fig. S13).

Our study adds *Drosophila* to the long list of insects that are attracted to CO_2_^38^. This implies an ancient evolutionary role of CO_2_ in insect behavior, as well as a highly conserved means for detecting it. These hypotheses are supported by our finding that CO_2_ attraction in *Drosophila* requires the ionotropic co-receptor IR25a, the most highly conserved olfactory receptor among insects^39^ (over 550-850 million years old^40^). Curiously, attraction to CO_2_ in mosquitoes (as well as members of Coleoptera (*Tribolium castaneum*) and Lepidoptera (*Bombyx mori*)) is mediated—at least in part—by a system homologous to the Gr63a/Gr21a gustatory receptors that mediate aversion in *Drosophila*^41^. Other insect species that respond to CO_2_, including members of Hymenoptera (honeybees^42^ and ants^12^), Hemiptera (bed bugs^3^ and kissing bugs^43^), Blattodea (termites^11^), and Ixodida (ticks^4^), however, lack this receptor^41^. It is possible that these insects also use the same evolutionarily ancient IR25a dependent CO_2_ pathway that is responsible for attraction in *Drosophila*.

The different time course in attraction to CO_2_ and ethanol, as well as the state-dependent decision to move towards or away from CO_2_, make this system ripe for exploring ecologically relevant decision making. Unfortunately, the GAL4 driver for the IR25a promoter is only expressed in about half of the endogenous IR25a-expressing neurons^44^, making imaging, silencing, and activation experiments difficult to interpret at this time. By narrowing the possible pathways of CO_2_ and ethanol attraction to IR25a, we hope to motivate future efforts to develop new genetic reagents that will make it possible to study this system in greater detail.

## Acknowledgements

We to thank Andrew Straw for providing the 3D tracking software for our flight experiments. Richard Benton offered helpful feedback on an early draft of the manuscript and also provided the IR8a; IR25a; Orco, Gr63a quadruple mutant. Ralf Stanewsky provided an IR25a mutant and the IR25a + BAC rescue line. Greg Suh provided an IR8a mutant. Elizabeth Hong and Jeff Riffell contributed helpful comments on the manuscript.

